## Supporting Information for "SMC complex unidirectionally translocates DNA by coupling segment capture with an asymmetric kleisin path"

### SUPPORTING TEXT

#### Validation of the reliability of all-atom force fields in the bottom-up coarse-graining approach.

In this study, we adopted a bottom-up approach to model hydrogen-bond-like interactions between protein and DNA interactions in a coarse-grained model. In this approach, the accuracy of all-atom simulations potentially influences the description of the coarse-grained model. In particular, the hydrogen bond patterns might vary depending on the choice of force field set used in the all-atom simulations.

To validate the robustness of the force field parameter set used in this study (Amber ff19SB for proteins (Tian et al., 2020), OL15 for DNA (Zgarbová et al., 2015), and OPC for water (Izadi et al., 2014)), we investigated force field dependency of the hydrogen bonding patterns between SMC ATPase heads and DNA.

In addition to the original parameter set (Amber ff19SB for proteins, OL15 for DNA, and OPC for water), we performed all-atom simulations in which the protein force field was changed to Amber ff99SB-ILDN (Lindorff-Larsen et al., 2010), the DNA force filed to Bsc1 (Ivani et al., 2016), and the water model to TIP4P-D (Piana et al., 2015). Here, we adopted Mg2+ ion parameters optimized for the TIP4P-D water model (Grotz and Schwierz, 2022). The result of the fraction of hydrogen-bond forming residues is shown in Fig. S2B. Although there were variations in the fractions and rankings, the top six amino acid residues that form hydrogen bonds with DNA are common between the original and the second force field parameter sets: 120Arg, 123Arg, 63Ile, 111 Arg, 62Arg, and 56Lys. Importantly, biochemical experiments have demonstrated the crucial roles of Arg62, Arg120, and Arg123 in the interaction between SMC ATPase heads and DNA (Vazquez Nunez et al., 2019). Even in amino acid residues with a lower fraction of hydrogen bond formation, we found common amino acid residues between two force field parameter sets such as 115Trp, 107Tyr 105Arg, 124Ser, 54Ser, 119Arg, and 112Ser and so on. Such consistency between two force field parameter sets supports the reliability of the original force field set (Amber ff19SB + OL15 + OPC).

Furthermore, we conducted all-atom simulations with a third set of force fields: Amber ff99SB-ILDN for proteins (Lindorff-Larsen et al., 2010), bsc1 for DNA (Ivani et al., 2016), and TIP3P for water (Jorgensen et al., 1983). As shown in Fig. S2C, we find that identified hydrogen-bond forming residues are essentially the same as in the original force field parameter set (Amber ff19SB + OL15 + OPC) and the second force field sets (Amber ff99SB-ILDN + Bsc1 + TIP4P-D).

These results clearly show that the choice of force field, water models, and magnesium ion parameters has only a minor impact on the hydrogen bond formation pattern between SMC ATPase and the DNA. Therefore, we conclude that the set of force fields we used in the bottom-up approach (*i.e.,* Amber ff19SB + OL15 + OPC water) is reliable for capturing the key amino acid residues and modeling coarse-grained hydrogen bond-like interactions.

### SUPPORTING FIGURES

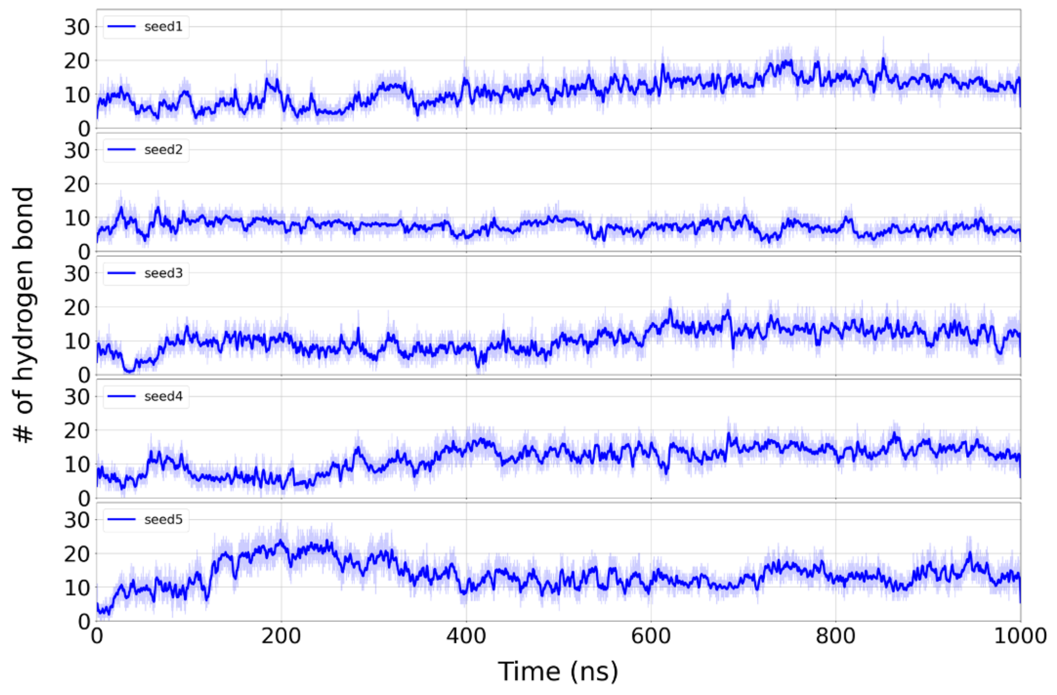


**Figure S1:** Time series of the number of hydrogen bonds between *Pf*SMC ATPase and DNA derived from all-atom MD simulations.

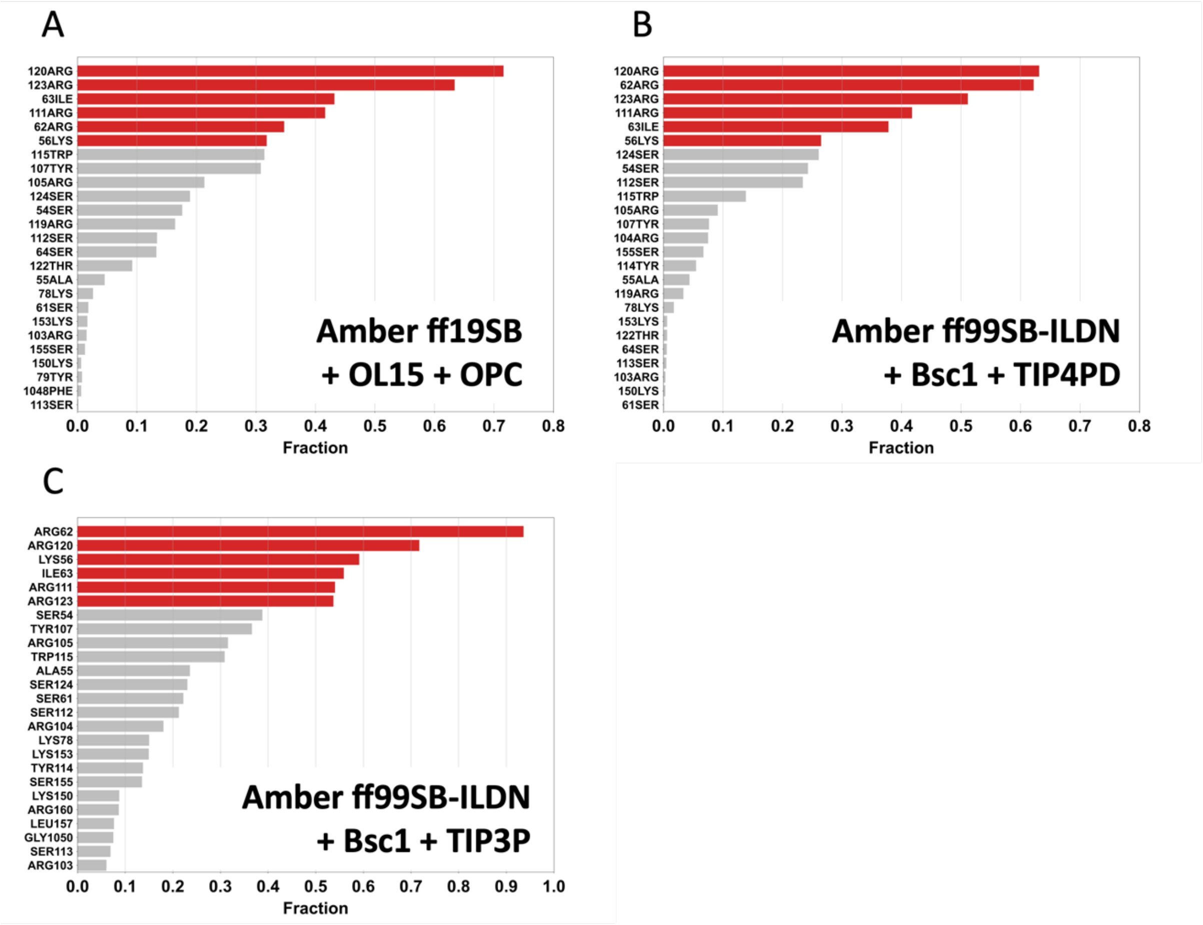


**Figure S2:** Force field dependence of tendency of hydrogen bond formation between SMC ATPase heads and DNA. (A) Fraction of hydrogen bond using Amber ff19SB for protein, OL15 for DNA, and OPC model for water. (B) Fraction of hydrogen bond using Amber ff99SB-ILDN for proteins, Bsc1 for DNA, and TIP4PD model for water. (C) Fraction of hydrogen bond using Amber ff99SB-ILDN for protein, Bsc1 for DNA, and TIP3P model for water.

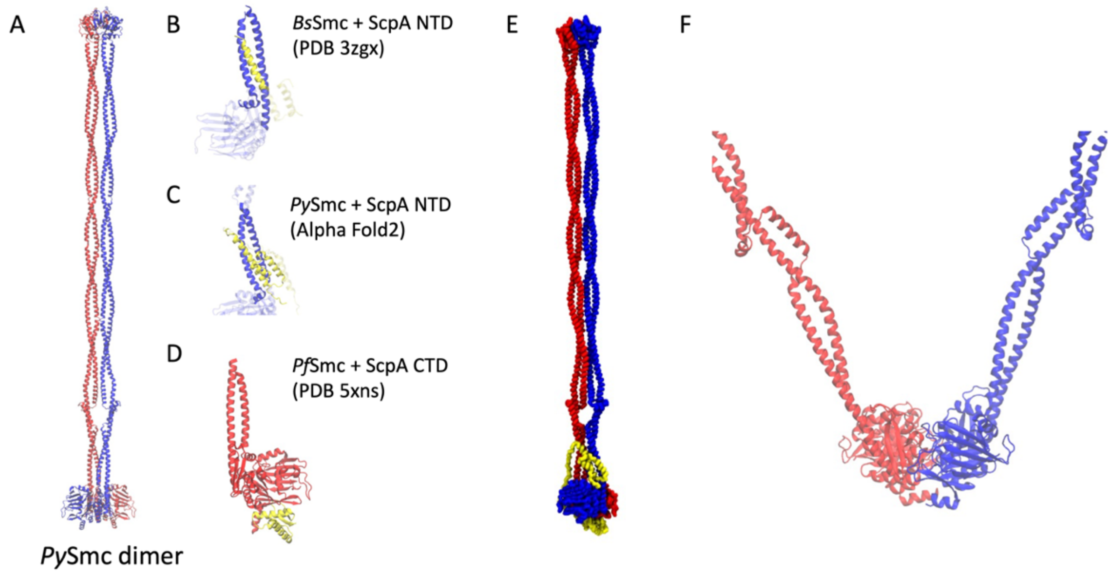


**Figure S3:** Modeling of a full-length *Py*SMC-ScpA complex. (A-D) Template structures for homology modeling of a full-length *Py*SMC-ScpA complex. The regions ignored in the homology modeling are indicated by transpare (E) A coarse-grained full-length *Py*SMC-ScpA complex model where one amino acid is represented by one bead. (F) A homology model of the engaged ATPase heads based on a crystal structure (PDB code: 1xex). The coiled-coil arms connected to the ATPase heads are modelled based on the I-shaped SMC dimer model in panel A.

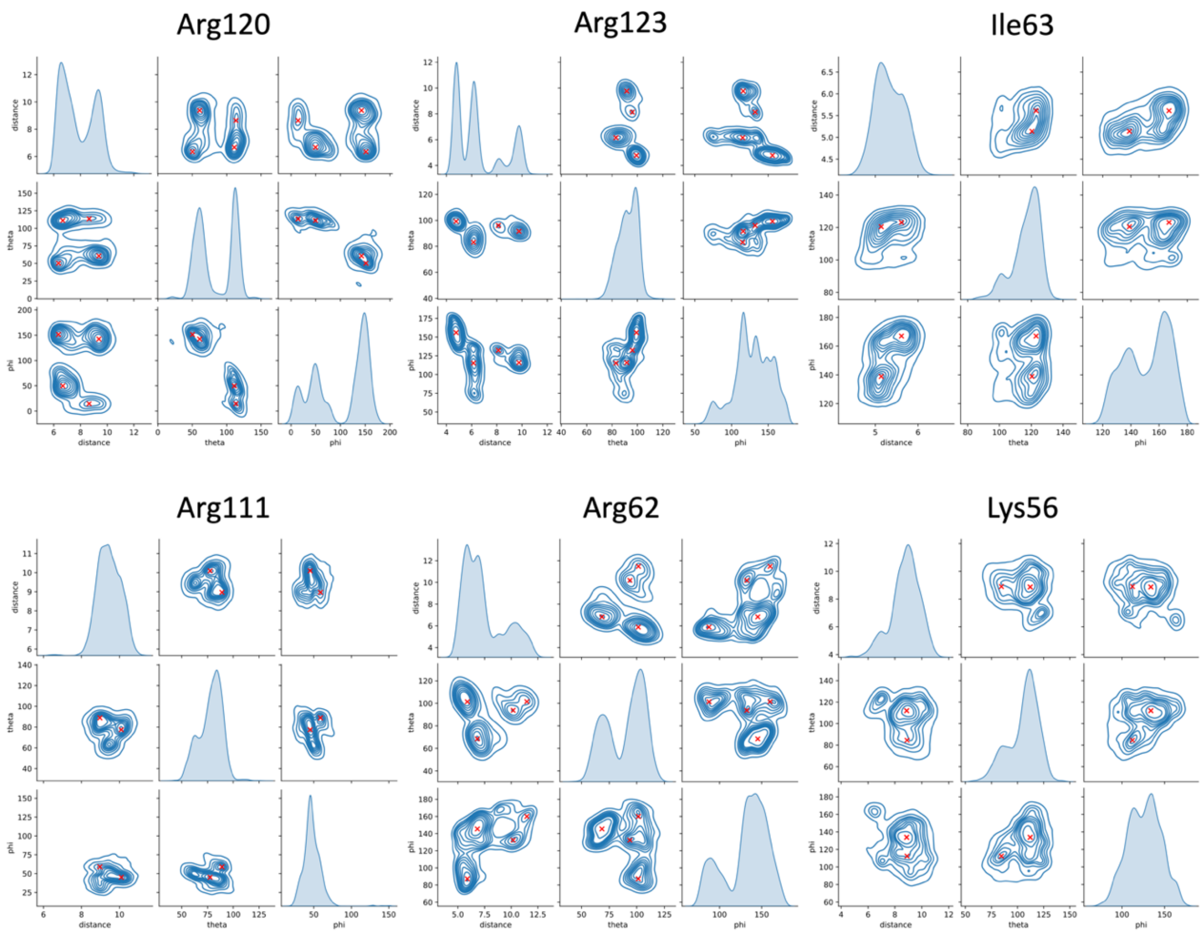


**Figure S4:** Probability distribution of the coarse-grained hydrogen bond parameters, $r$, $\theta$, and $\varphi$ for frequently hydrogen-bond forming residues. The red points are optimized hydrogen-bond parameters, which corresponding to peak positions.

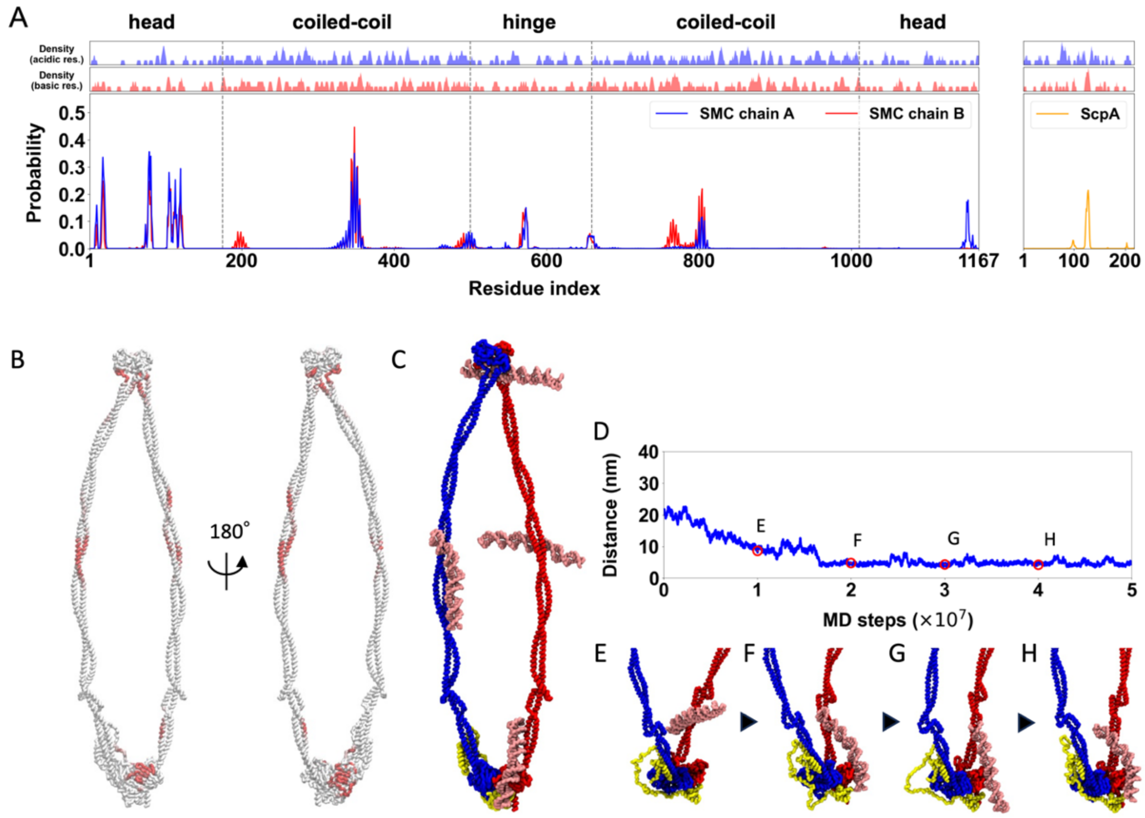


**Figure S5:** DNA binding sites in the SMC-ScpA complex where hydrogen bond interactions on the ATPase heads are not incorporated. (A) Top two panels plot the local average of charges defined as the moving average with window size of 5 residues. Bottom panel plots the contact probability between DNA and SMC-ScpA complex. (B) DNA contact probability mapped on the SMC-ScpA structure. (C) A typical snapshot of DNA binding to the SMC-ScpA complex. The DNA that binds to the ATPase heads does not migrate to the top, staying on the side surface of the ATPase heads. (D) A representative timeseries of the distance between center of mass of the ATPase heads and DNA. (E-H) Representative snapshots during a DNA binding event to the SMC ATPase heads.

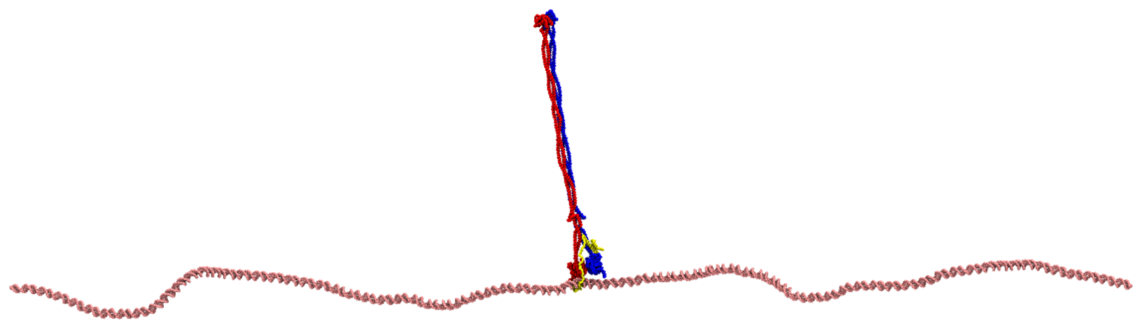

**Figure S6:** An initial structure of DNA translocation simulations. An 800 bp dsDNA was placed into the kleisin ring of the disengaged state of the SMC-ScpA complex.

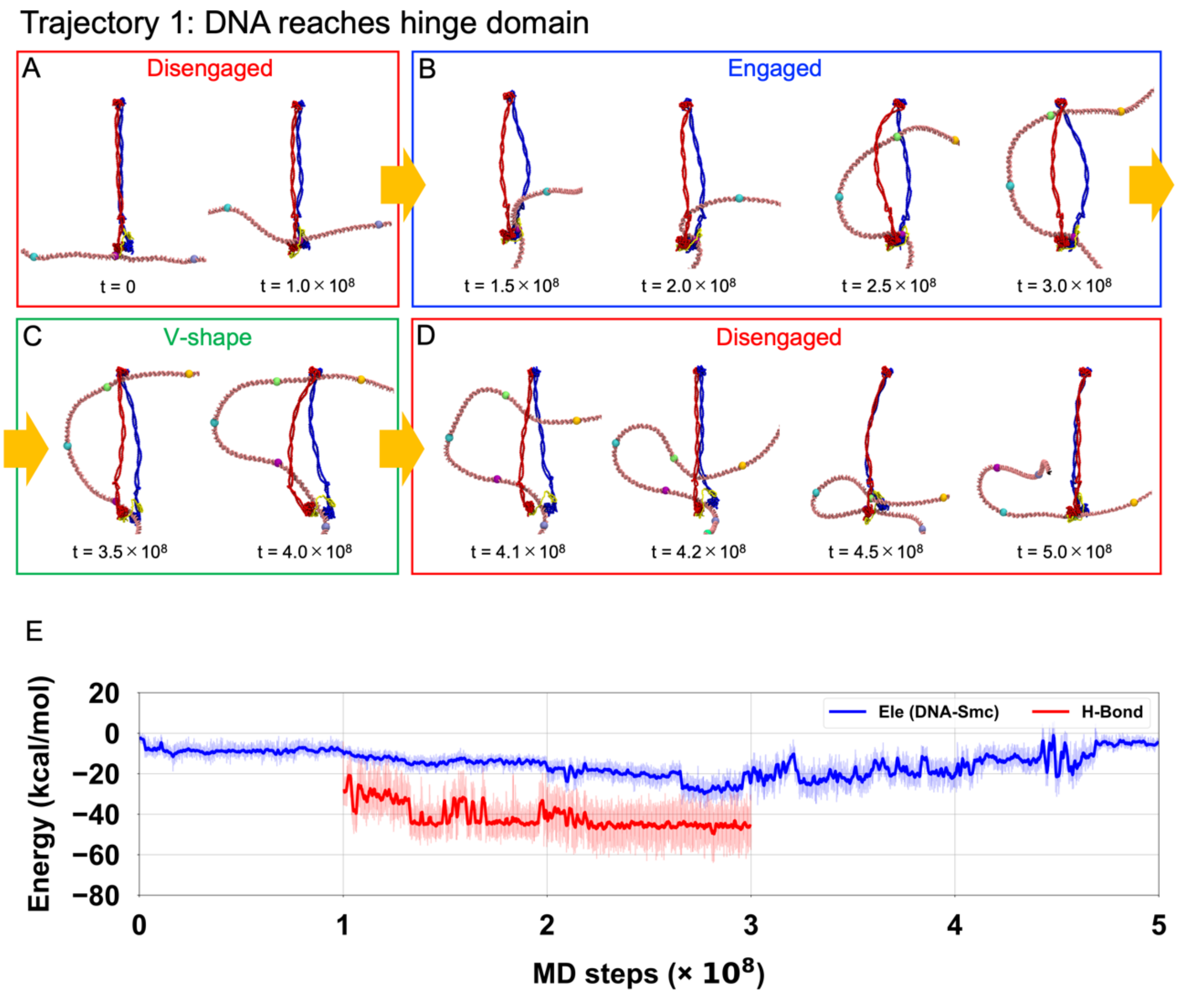


**Figure S7:** Detail snapshots during DNA translocation via DNA-segment capture. (A)The disengaged state during $t = 0$ - $1.0{\times}10^8$ MD steps. (B) The engaged state during $t = 1.0{\times}10^8$ - $3.0{\times}10^8$ MD steps. (C) The V-shape state during $t = 3.0{\times}10^8$ - $4.0{\times}10^8$ MD steps. (D) The disengaged state $t = 4.0{\times}10^8$ – $5.0{\times}10^8$ MD steps. (E) Tims series of electrostatic and hydrogen bonding interactions between the SMC complex and DNA.

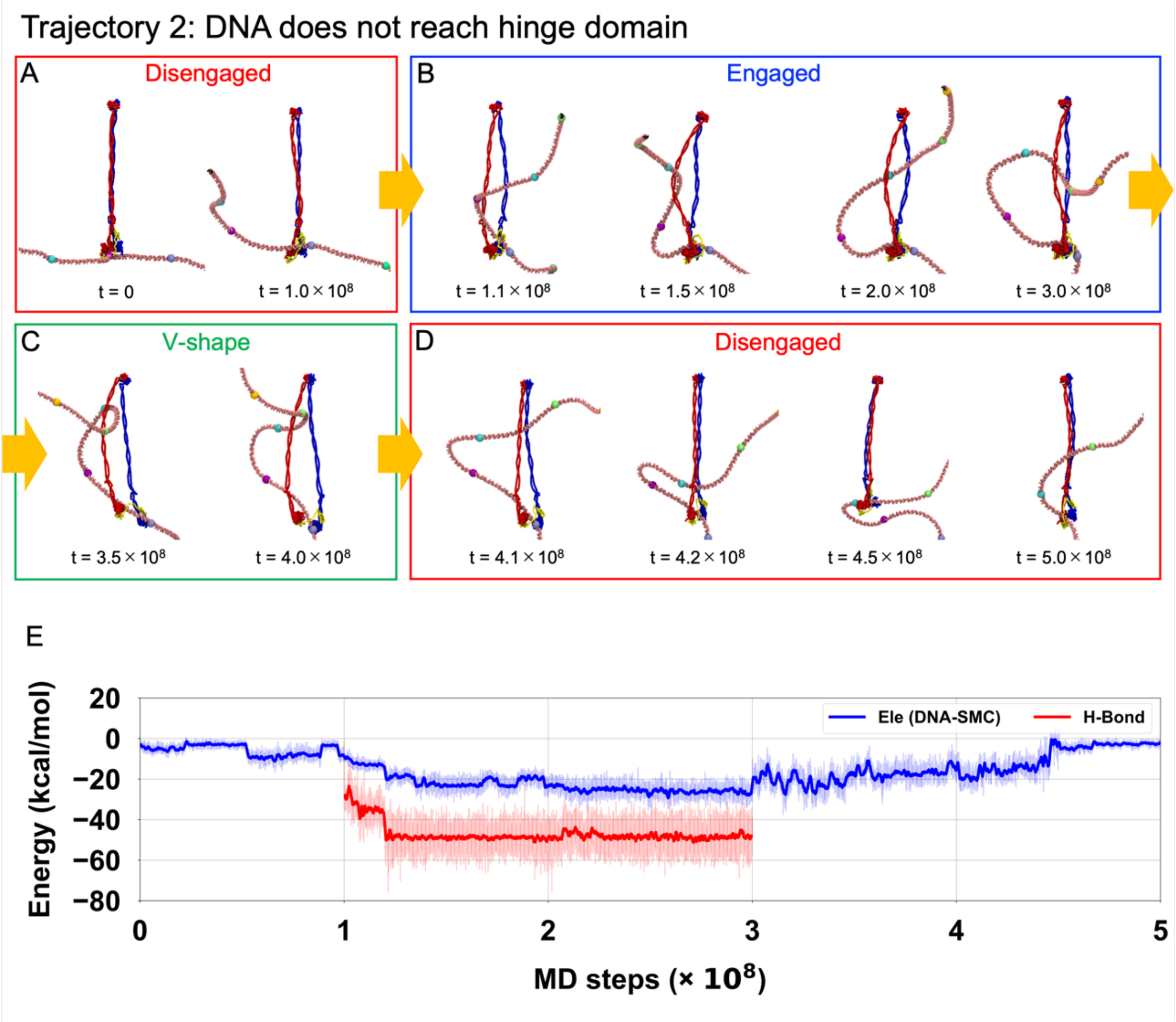


**Figure S8:** Detail snapshots during DNA translocation via DNA-segment capture where the DNA does not reach the hinge domain in the engaged state. (A)The disengaged state during $t$ = 0 - 1.0×$10^8$ MD steps. (B) The engaged state during $t$ = 1.0×$10^8$ - 3.0×$10^8$ MD steps. (C) The V-shape state during $t$ = 3.0×$10^8$ - 4.0×$10^8$ MD steps. (D) The disengaged state $t$ = 4.0×$10^8$ – 5.0×$10^8$ MD steps. (E) Tims series of electrostatic and hydrogen bonding interactions between the SMC complex and DNA.

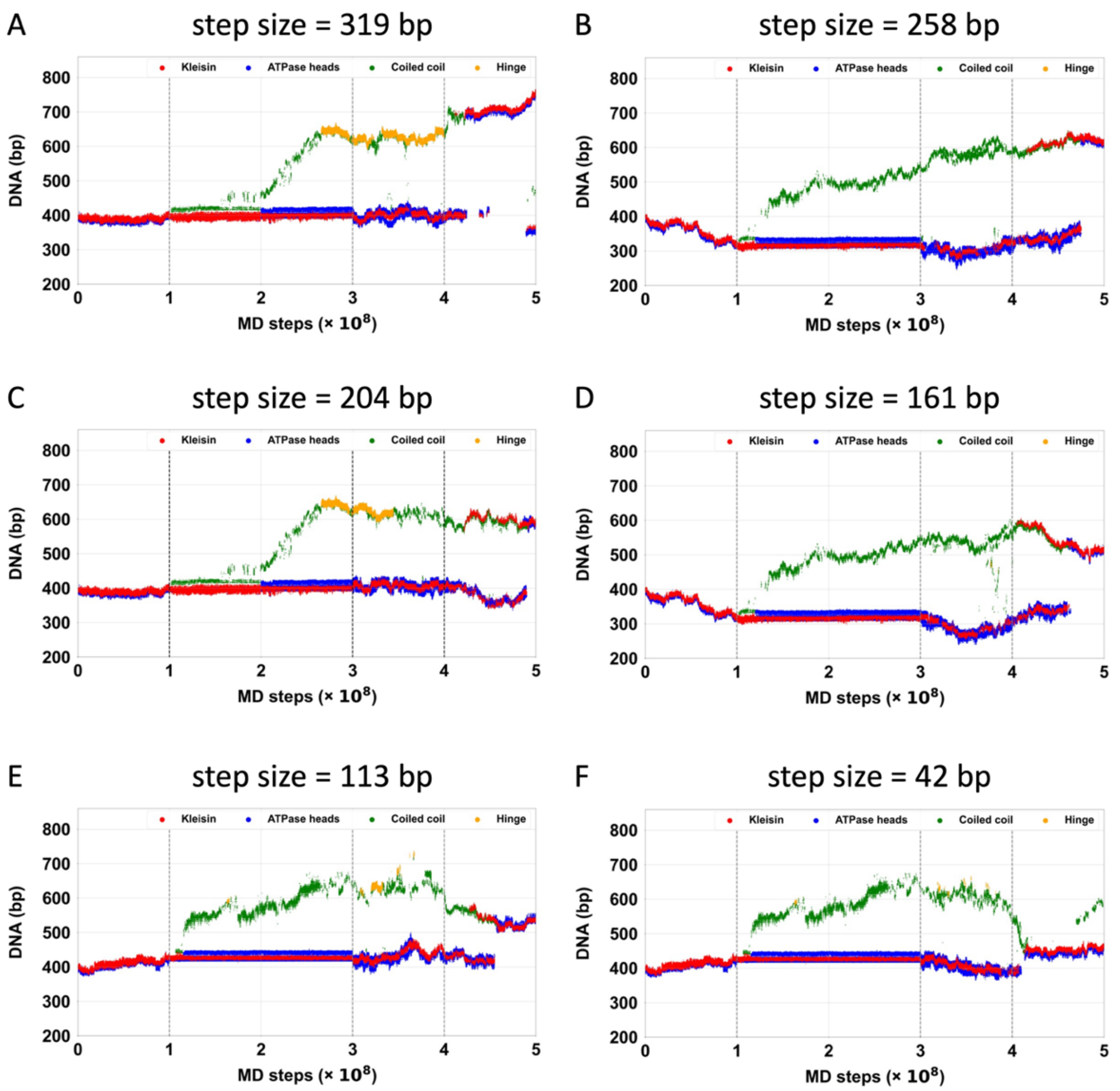


**Figure S9:** Time series of the DNA position where each SMC and kleisin domain contacts. DNA-Protein contacts at kleisin, ATPase heads, coiled-coil, and hinge domains are plotted in red, blue, green, and orange, respectively. SMC-ScpA complex realizes a variety of step sizes such as (A) 310 bp, (B) 258 bp, (C) 204 bp, (D) 161 bp, (E)113 bp, and (F) 42 bp.

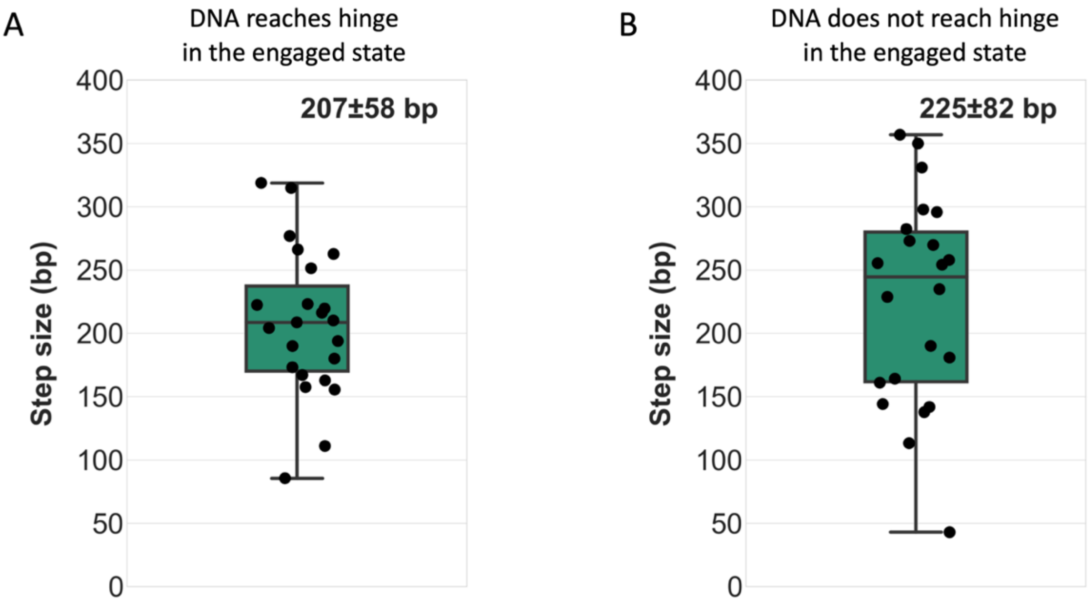


**Figure S10:** Translocation step size for (A) trajectories in which the DNA reaches the hinge domain and for (B) trajectories in which the DNA does not reach the hinge in the engaged state.

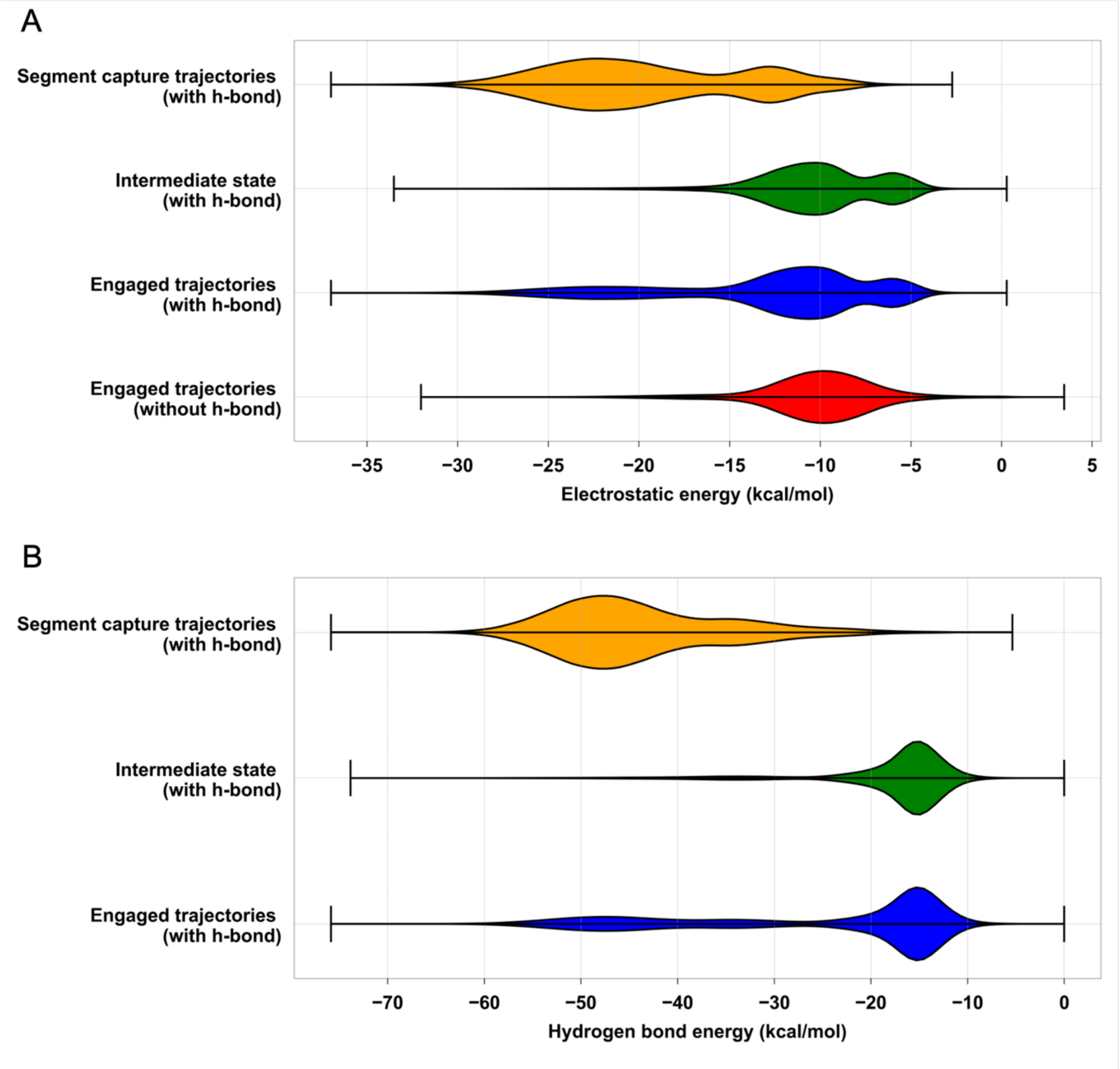


**Figure S11:** Probability distribution of (A) electrostatic and (B) hydrogen bonding energies between the SMC complex and DNA.

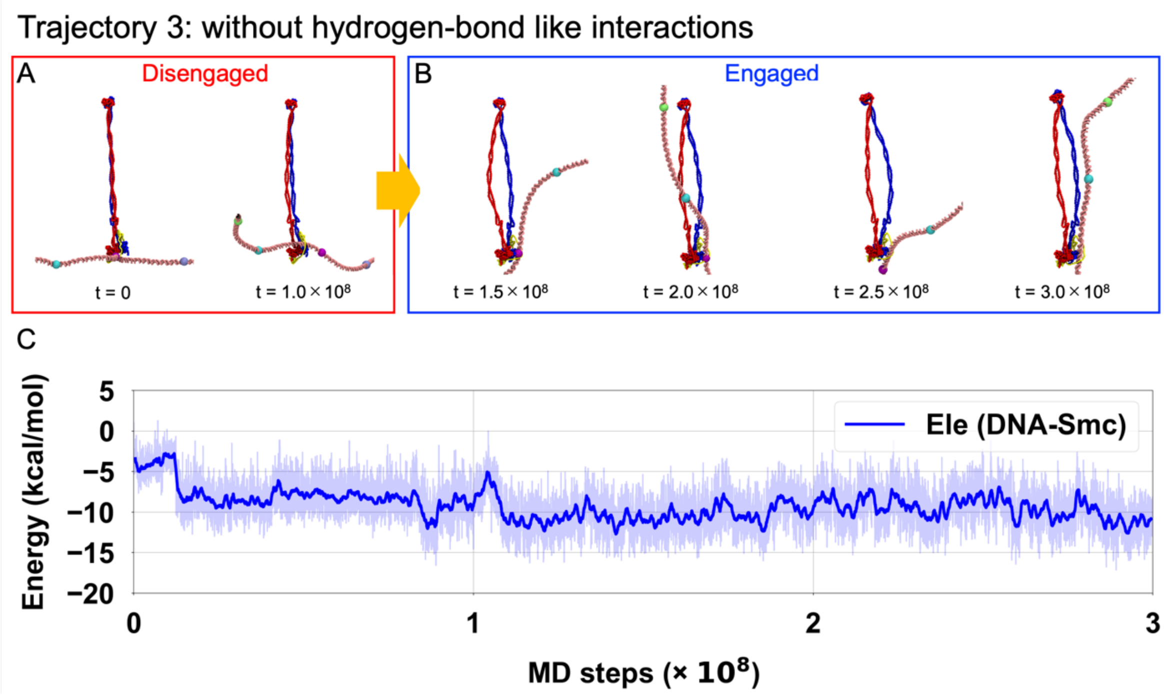


**Figure S12:** Detail trajectories of simulations for SMC complex and DNA without hydrogen bonding interactions between SMC ATPase heads and DNA (A)The disengaged state during $t$ = 0 - 1.0×$10^8$ MD steps. (B) The engaged state during $t$ = 1.0×$10^8$ - 3.0×$10^8$ MD steps. (C) Tims series of electrostatic interactions between the SMC complex and DNA.

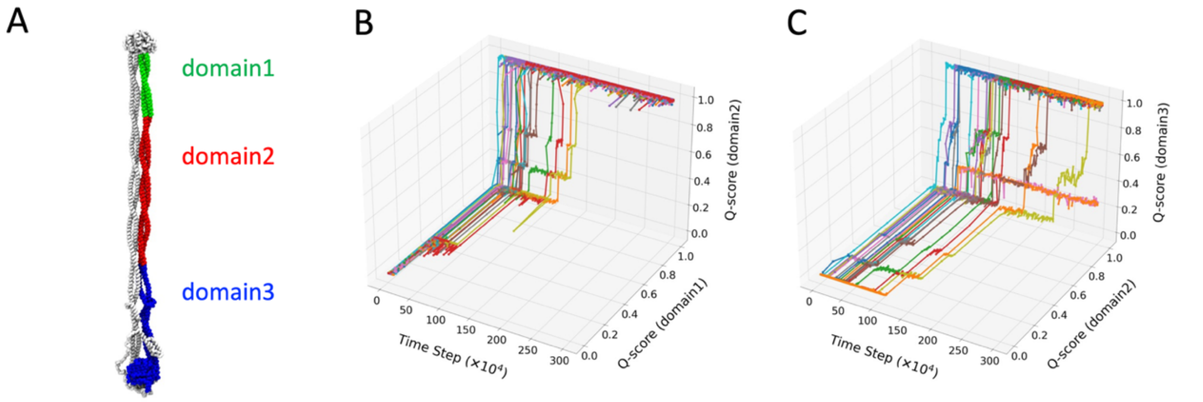


**Figure S13:** Zipping up the coiled-coil arms when transition from V-shape to disengaged states. (A) The definition domains where the coiled-coil arm and ATPase heads are divided into three domains. (B) Time series of Q scores for domain 1 and domain 2 for all simulations. (B) Time series of Q scores for domain 2 and domain 3 for all simulations.

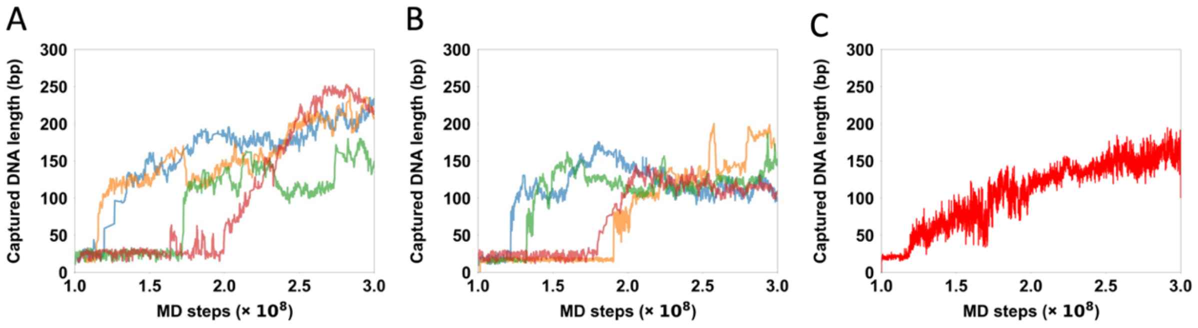


**Figure S14:** Time series of the captured DNA length within the SMC ring in the engaged state during $t = 1.0\times10^8 - 3.0\times10^8$ MD steps. (A) Captured DNA length for the trajectories that lead to the DNA translocation successfully in the subsequent states. (B) Captured DNA length for the trajectories that do not lead to the DNA translocation in the subsequent states. (C) Time series of the averaged captured DNA length.

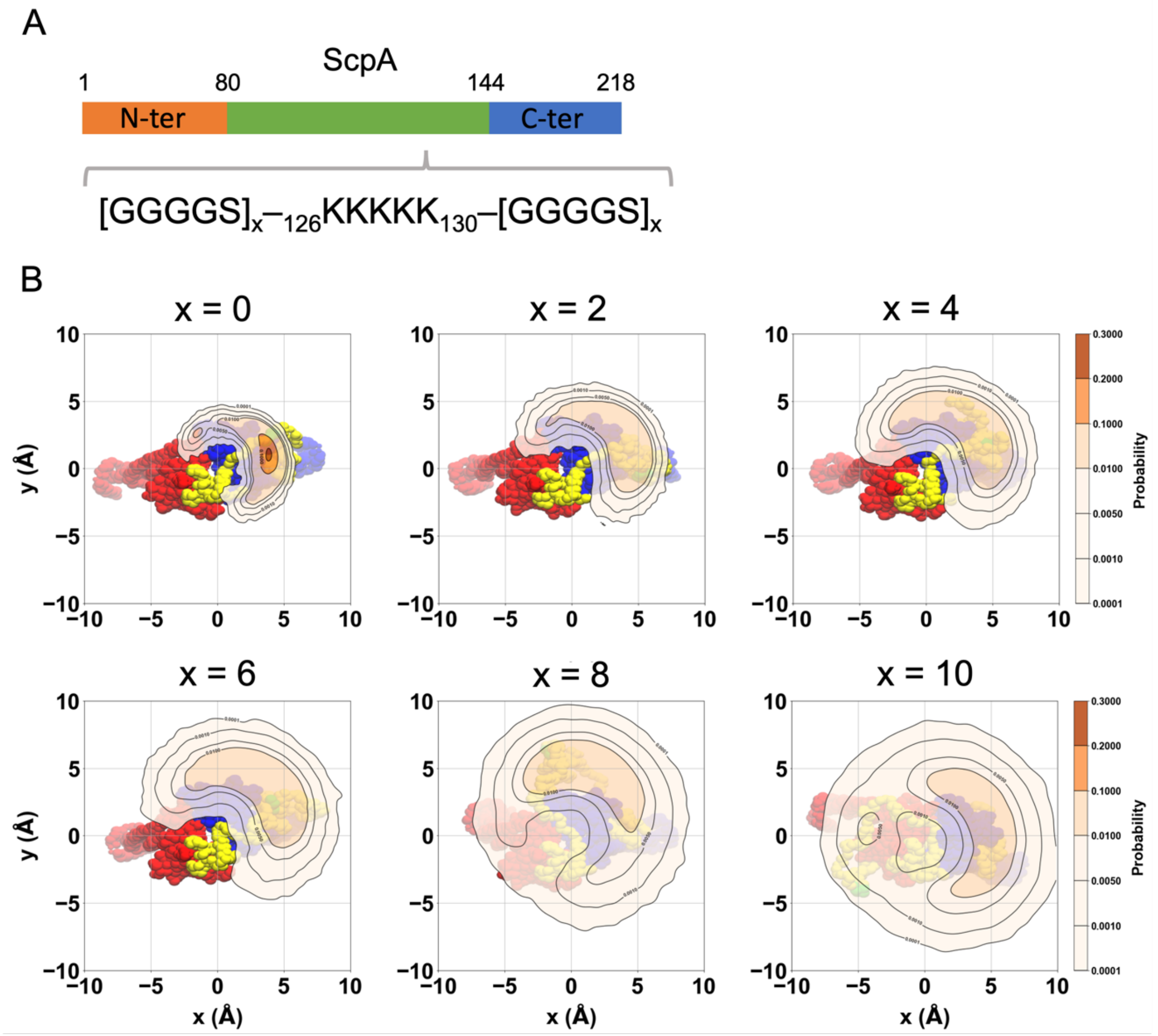


**Figure S15:** The spatial distribution of the DNA patch on the ScpA subunit that have different linker length. (A) A schematic figure showing the position of the GGGGS linker introduced into the ScpA subunit. (B) The spatial distribution of the DNA patch.
